# Comparison of manual and automated ventricle segmentation in the maternal immune stimulation rat model of schizophrenia

**DOI:** 10.1101/2020.06.10.144022

**Authors:** Rebecca Winter, Benson Akinola, Elizabeth Barroeta-Hlusicka, Sebastian Meister, Jens Pietzsch, Christine Winter, Nadine Bernhardt

## Abstract

Maternal immune stimulation (MIS) is strongly implicated in the etiology of neuropsychiatric disorders. Magnetic resonance imaging (MRI) studies provide evidence for brain structural abnormalities in rodents following prenatal exposure to MIS. Reported volumetric changes in adult MIS offspring comprise among others larger ventricular volumes, consistent with alterations found in patients with schizophrenia. Linking rodent models of MIS with non-invasive small animal neuroimaging modalities thus represents a powerful tool for the investigation of structural endophenotypes. Traditionally manual segmentation of regions-of-interest, which is laborious and prone to low intra- and inter-rater reliability, was employed for data analysis. Recently automated analysis platforms in rodent disease models are emerging. However, none of these has been found to reliably detect ventricular volume changes in MIS nor directly compared manual and automated data analysis strategies. The present study was thus conducted to establish an automated, structural analysis method focused on lateral ventricle segmentation. It was applied to ex-vivo rat brain MRI scans. Performance was validated for phenotype induction following MIS and preventive treatment data and compared to manual segmentation. In conclusion, we present an automated analysis platform to investigate ventricular volume alterations in rodent models thereby encouraging their preclinical use in the search for new urgently needed treatments.

## Introduction

Exploring new avenues in psychiatric research has the goal of combing treatment effectiveness with tolerability, often with the idea of a preventive treatment strategy. One condition in order to achieve this high-set goal is the availability of cross-species biomarkers. Good parameters should be easily measured, reliably report disease manifestation in patients and inform preclinical treatment studies in disease relevant animal models. Discovering biomarkers has proven to be very challenging for many psychiatric diseases, due to the variability of symptoms and their often late onset. Most psychiatric diseases, like schizophrenia, are now considered neurodevelopmental, which means that the pathological alterations occur during development and are present before the symptoms develop. These pathological alterations can for example be seen in morphological changes in the brain, e.g. divergent volume of brain regions. One of these locations are the lateral ventricles, which are altered in a number of psychiatric or neurological afflictions: for example in ischemic stroke patients (1), amyotrophic lateral sclerosis (2), cystic periventricular leukomalacia (3), Alzheimer’s disease (4), Parkinson’s disease (5) and schizophrenia (6,7). Similar changes in lateral ventricle volumes have been observed in a number of established animal models with relevance to schizophrenia (8–10) as well as other psychiatric or neurological diseases (11–15).

Brain volumes can be easily measured with magnetic resonance imaging (MRI), a non-invasive imaging method available with a good soft-tissue contrast and good spatial resolution (50 - 100 µm) (16,17). It utilizes longitudinal magnetization signals to create contrast images whose intensity range indicate structural boundaries and tissue properties in order to further make informed decisions. To quantify structural alterations due to neuropsychiatric diseases, MR images are acquired and morphometry is performed. Several morphometric methods have been established, including voxel based morphometry (VBM) (18) to correlate brain structure or shape differences within a population, cortical thickness measurement (19,20), statistical shape analysis (21) and structural volume analysis (22) with labeling of anatomical structures. For the process of morphometric analyzing the structural MR images for volume, shape and/or thickness, segmentation of the volume images is typically carried out. Segmentation is a process of classification of image units (pixels or voxels) into subsets based on predefined criteria. The manual segmentation of the rodent’s brain is often still the standard for segmentation of small animal structural images (23). This method is very laborious, time consuming and subjected to intra- and inter-subject variability. In the search of urgently needed new treatment strategies however, there is a high demand for reliable automated processing alternatives (24,25).

For automated morphometric analysis of the rodent brain it is important to accurately determine the anatomical structural region’s parcellation. Any established method must be reliable and allow repetitiveness. A method that fits this need is the atlas-based segmentation. The automated atlas-based segmentation of volumetric images is based on image registrations and label propagation. It involves the registration (linear and diffeomorphic) between a reference brain image to individuals of a query set (or vice versa). The resulting registered brain represents the maximum similarity between the individual image and the corresponding reference image. The anatomical segmentation or label of the reference brain is further propagated to the individual brain sets based on the transformations and deformation fields from the registration step. Registration techniques, variation between animals’ anatomical structures and correctness of label segmentation are critical factors for the automated segmentation of the rodent brain using an atlas-based technique. More recently, some more automated workflows have been implemented (26,27). These approaches are less labor-intensive and time-consuming; however, they cannot be directly applied to our data depending on differences between species (mouse vs rat), strains, gender or age factors. In addition, especially segmentation of lateral ventricles has been proven difficult (26).

Thus, we aimed to establish an automated analysis platform with a focus on lateral ventricle delineation. For this we chose a maternal immune stimulation (MIS) rat model for schizophrenia, which is based on the finding that maternal prenatal infection increases the risk of developing schizophrenia (Brown et al, 2004). Exposing pregnant rodents to the viral mimic polyriboinosinic polyribocytidilic acid (poly I:C) is a commonly used neurodevelopmental approach to model schizophrenia (8–10,28,29). The onset of behavioral abnormalities is seen during adolescence, similar to the clinical picture in humans (10). However, the neuropathological alterations have been detected at a different time point with the enlargement of lateral ventricles occurring in adulthood (30). This allows studies of preventive treatment approaches. In line, we have previously reported that anodal transcranial direct current stimulation (tDCS) has been shown to prevent the emergence of behavioral alterations and the enlargement of lateral ventricles in the MIS animal model (8). Here, we report an analysis platform construction optimized for lateral ventricle volume estimation in adult male offspring following prenatal MIS induction. In addition, MR images were segmented manually and obtained results compared between methods. Finally, available imaging data from our prior tDCS treatment study was employed for verification of platform performance and data robustness.

## Materials and methods

### Animals

Experiments were performed according to the guidelines of the European Union Council Directive 2010/63/EU for care of laboratory animals approved by the local ethic committee (Landesdirektion Sachsen, Dresden, Germany). Animals were housed in a temperature and humidity-controlled vivarium under a 12 h light / 12 h dark standard day cycle with food and water ad libitum. Animal suffering and numbers were kept to a minimum. Female pregnant Wistar rats (Charles River Laboratories, Europe) were received at gestational day (GD) 15, housed in individual cages with nesting material and left to acclimatize for 2 hours. Dams were carefully anesthetized with isoflurane and the viral analogue poly I:C, (4 mg/kg; Sigma, Germany) dissolved in saline, or 0.9% saline alone was injected through the tail vein (volume: 100 µl/100 gr bodyweight) (31,32). Dams were weighed daily until delivery and then left undisturbed until postnatal day (PND) 21, when offspring was weaned, group-housed with 2-4 animals in standard cages with bedding and shelter (dark PVC tubes). Adult (>PND90) male offspring, n=21 from saline and n=20 from poly I:C injected mothers, were used for the study.

Second, imaging data from a previous published study (8) was utilized. The study was conducted to assess the preventive treatment potential of non-invasive transcranial direct current stimulation (tDCS) during adolescence, prior to schizophrenia-relevant behavioral manifestation, in the same MIS model of schizophrenia. The study followed a 2 × 3 design with phenotype (saline, poly) × treatment (sham, cathodal, anodal), for details refer to Hadar et al., 2019. Groups: saline-sham (ss) n = 15, poly-sham (ps) n = 14, saline-cathodal (sc) n = 13, poly-cathodal (pc) n = 11, saline-anodal (sa) n = 11 and poly-anodal (pa) n = 12.

### MRI

#### Sample preparation

For postmortem assessments the rats were deeply anaesthetized with a single i.p. injection of pentobarbital (60 mg/kg) and perfused transcardially with cold 4% paraformaldehyde in 0.1 M phosphate buffer, pH 7.4. Brains remained within the scull and were imaged within 1-2 weeks following preparation.

#### Acquisition

MRI acquisitions were performed on a 7 Tesla rodent scanner (Bruker BioSpin MRI GmbH, Ettlingen, Germany) at Charité (Berlin, Germany) or HZDR (Dresden-Rossendorf, Germany). The acquisition protocol consisted of a multi slice localizer (field of view (FOV) 50 × 50 mm) and T2-weighted contrast images with a rapid acquisition with relaxation enhancement (RARE) sequence (imaging parameters: TR/TE = 4050/30 ms, RARE factor 8, NEX 6, FOV 30 × 30 mm, MD 256 × 256), resulting in in an inplane resolution of 117 *µ*m with a slice thickness of 0.5 mm and 42 slices. Scanning time was 13 minutes per brain.

#### Manual segmentation

The axial view was used for manual segmentation. Original acquired images were resliced with a slice thickness of 0.12 mm. 15 slices from −0.1mm (first view of the anterior part of the anterior commissure) to +0.7mm from Bregma were analyzed for each animal (33). For best image quality, contrast was adjusted using MRIcron. The lateral ventricles were outlined using ImageJ and special attention was paid on distinguishing ventricular area from adjoining areas through identifying differences in contrast by eye. Manual segmentation took up to 120 minutes per animal, as both left and right lateral ventricle were separately measured through each slice and discernable contrasts were often minimal, which required extensive repeated examination. Two independent raters carried out manual segmentation. Volumes were obtained by multiplying the summed-up amount of pixels times pixel area with slice thickness (ventricular volume = (sum (measured amount of pixels × 0.117mm^2^)) × 0.12mm × 0.01). Analysis were run on means from the two independent observations per animal.

#### Automated analysis platform

The DICOM (Digital Imaging and Communications in Medicine) MR image data were collected, converted into the Neuroimaging Informatics Technology Initiative (NIFTI) file format and utilized for the platform’s framework and evaluation. To provide computational efficiency the automated analysis platform was based on the Advanced Normalization Tools (ANTs) (34), a toolkit for medical image registration and segmentation and was implemented with a high-performance computing (HPC) setup (see S1 Appendix for details). The structure of the automated analysis platform is shown in Fig 1.

**Fig 1.**
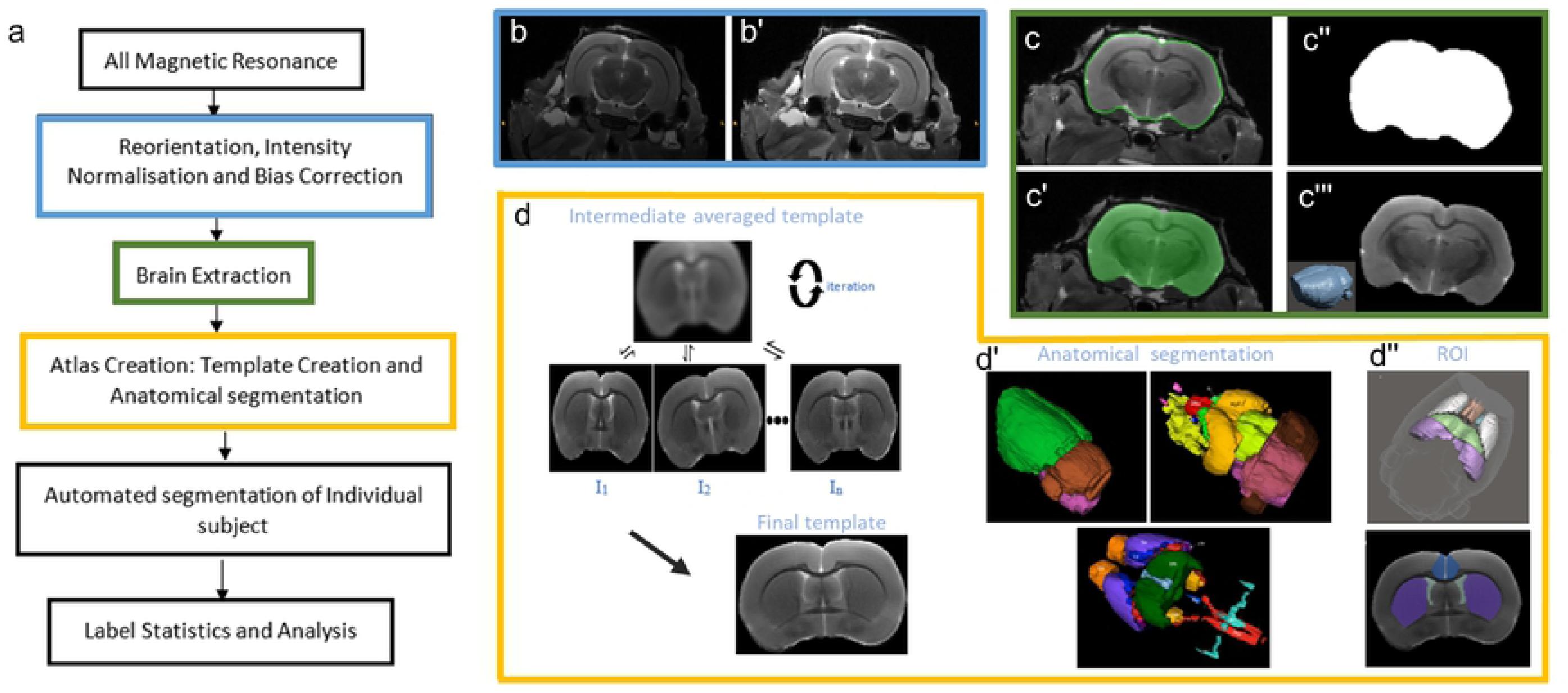
Automated analysis platform overview. (a) Flowchart with examples of b) bias correction including intensity normalisation followed by field correction for inhomogeneity using the N4BiasFieldCorrection (64). c) Brain extraction with registration-based skull stripping to remove non brain tissues using a mask and the skullstripper tool (65). d) Atlas Creation: Iterative creation of the within study template using the minimum deformation pairwise registration resulted in a study template, an averaged image of selected control subjects d’ with its corresponding manually delineated labels and d’’ regions of interest (ROI) for studies in the maternal immune activation model.

Pre-processing steps included intensity reorientation, intensity bias correction and brain extraction. To achieve spatial consistency images were reoriented into standard anatomical space (patient coordinate) based on the ITK-SNAP (35) employing the *orient* algorithm of the *Convert3D* medical image processing tool. Hence data sets with an initial orientation LPI (Left, Posterior, Inferior), common for DICOM images i.e. an image vector were converted by a linear transformation to RIP (Right, Inferior, Posterior).

The N4BiasFieldCorrection algorithm (N4ITK), an optimized version of the N3 that implements a robust B-spline approximation routine and a modified optimization scheme (36), was used to correct intensity non-uniformity within images. 50 iterations over three levels with a convergence threshold of 1e-6, full width at half maximum (deconvolution kernel) of 0.15mm were employed with default parameters to provide optimum results for the rat subject MR images. For brain extraction the python command line *SkullStrip* (37) was used for the semi-automated registration brain extraction in this project. A reference stripped image with its brain mask (manually delineated) and the brain image was used to generate a stripped fixed image mask that was then binarized. Image multiplication operation was then performed with this mask and the original subject image (see S1 Appendix for details).

Post-processing steps included the segmentation of each subject, i.e. the image registration, normalization, label propagation and volume extraction. For the segmentation of each subject the atlas-based method was implemented using a within study created atlas from four control subjects (38). The *antsMultivariateTemplateConstruction.sh* script (39) of the ANTs tool was utilized for template creation. An iteration of four was employed with the greedy symmetric normalization (SyN) transformation with a cross correlation similarity metric (39,40). The anatomical image of the template was manually delineated into thirty-five regions using the ITK-SNAP. The standard rat brain atlas (33) was employed for anatomical referencing and nomenclature. The left and right hemispheres were considered as one unit. The ependymal and subependymal layers were considered the boundaries of the lateral ventricles (LV). Similarly, the periventricular hypothalamic region was classified with the 3rd ventricles.

The *antsRegistration* command (41) of the ANTs was employed for image registration. The cross-correlation (CC) similarity metric was used for diffeomorphic registration and the mutual information during linear registration (26,39,42) between the fixed and moving images. The symmetric normalization (SyN) (42) was the optimizer (image normalization method) algorithm used during deformation with the CC similarity (39). The antsApplyTransforms command of the ANTs tool was employed for label propagation of all subjects with a nearest neighbor interpolation scheme. From the automated segmentation description using an atlas-based segmentation technique, the quality of segmentation of the output is directly proportional to the performance of registration (43) and interpolation scheme (see S1 Appendix for details). The volume of each structure of the label was extracted using the LabelStats algorithm of ANTs and a MATLAB script.

### Statistics

One way ANOVA was used to determine significant differences between phenotype (saline or poly) for the lateral ventricle volumes measured through manual segmentation and the automated analysis platform, respectively. To check for significant interactions between phenotype (saline or poly) and treatment (saline, anodal or cathodal), two-way ANOVA was completed followed by posthoc analysis with Sidak correction. Statistical significance was set at p<0.05. The software SPSS was used for statistical analysis of the data.

## Results

MR images from 42 animals (n=21 saline, n=20 poly) were analyzed using the established analysis platform, labels were visually quality controlled but not corrected and ventricular volumes estimated. As expected, estimates were observed to be lower in saline 4.198 ± 0.247 mm^3^ compared to poly I:C 4.434 ± 0.279 mm^3^ offspring, respectively (Fig. 2a,b). The difference was found to be significant T(39)= −2.858 p= .007 (Fig 2c). Due to the enormous labor intensity only half of the sample was also evaluated manually (n=12 saline, n=12 poly; for example of manual segmented image see S1 Appendix). Analysis showed similarly lower estimations of LV volumes with 3.981 ± 0.463 mm^3^ and 4.293 ± 0.383 mm^3^ in saline compared to poly I:C offspring which did not reach significance T(22)= - 1.794 p= .087 (Fig. 2d). Reduced samples size and variance between samples in manual estimates, which is two times higher compared to platform measures, can be taken in consideration, however automated analysis platform data from an identically reduced sample remains significant T(22)= −2.481 p= .023. Thus both evaluation methods yield similar phenotype differences while the correlation between measures is small r(22)=.249, p=.032.

**Fig 2.**
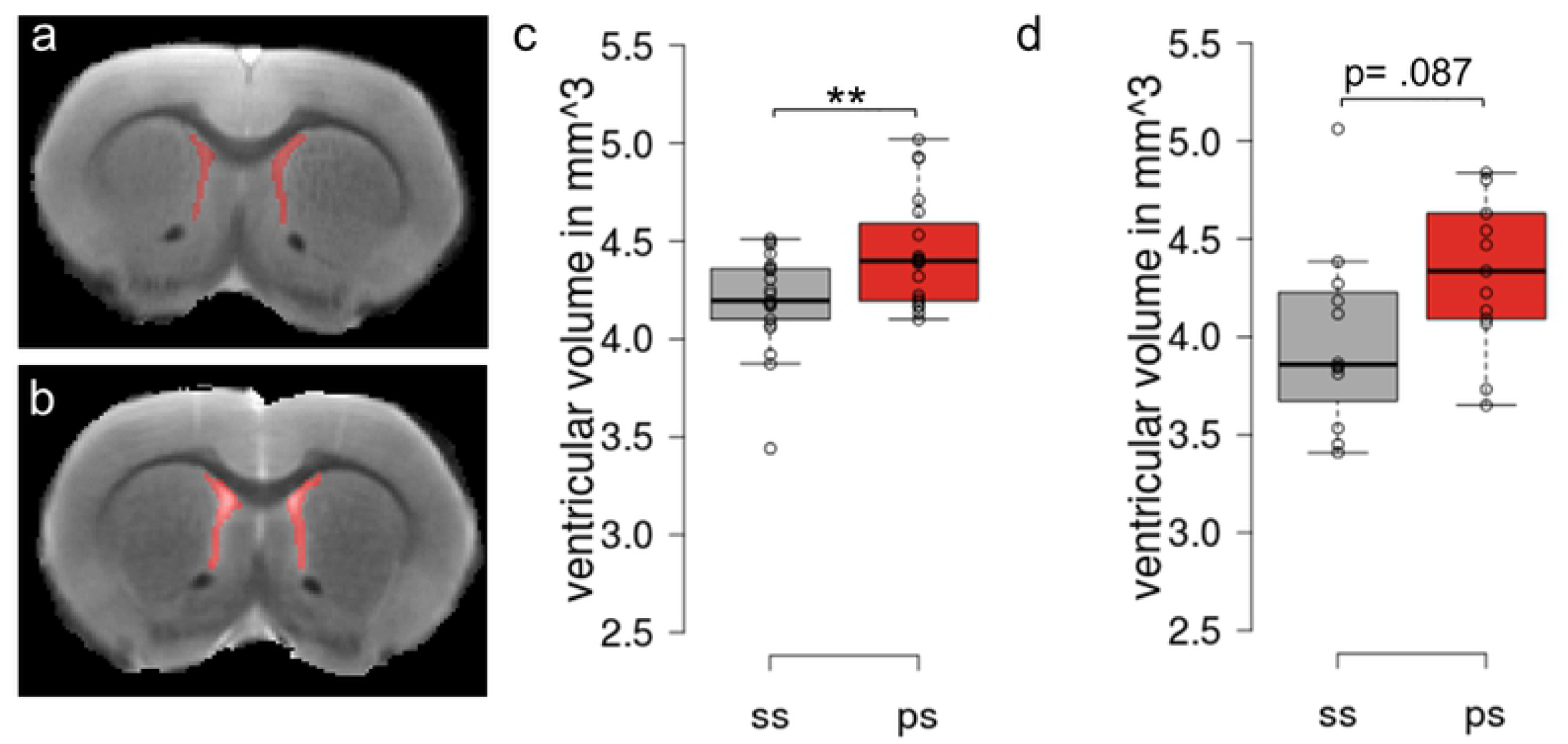
Lateral ventricle (LV) phenotype in rat MIS model. a,b) Examples of LV delineated from a saline a) and poly (b) subject by the automated analysis platform. Boxplot of the lateral ventricular volume from poly I:C and saline injected offspring following c) automated and d) manual delineation methods. Center lines show the medians; box limits indicate the 25th and 75th percentiles as determined by R software; whiskers extend 1.5 times the interquartile range from the 25th and 75th percentiles, individual data points included.

For further performance validation, imaging data from (8), which was published employing manual delineation, was now re-analyzed with the newly established automated analysis platform. Following visual quality control, ventricle labels were manually corrected for 15% of subjects and LV volume statistics obtained (Fig. 3). A two way ANOVA yielded a significant effect for the phenotype*treatment interaction (F(5,68) = 8.626, p < .001) with no main effects of phenotype or treatment (p > .05). Posthoc analysis showed the expected, significant phenotype difference following MIS between the saline control (ss) and the untreated poly I:C (ps) group p = .002, mean ventricular volume ss: 3.97 ± 0.25 mm^3^ and ps: 4.28 ± 0.21 mm^3^. Similarly, as measured and statistically evaluated in the Hadar et al., 2019 paper, treatment was found to reduce LV volume following MIS when compared to ps animals for cathodal p= .004 and anodal p=.006 stimulation, respectively. Mean ventricular volume for pc: 3.98 ± 0.18 mm^3^ and pa: 4.00 ± 0.30 mm^3^ groups. Finally, also differences between sa and pa were replicated p=.037, mean ventricular volume for sa: 4.2 ± 0.09 mm^3^.

**Fig. 3.**
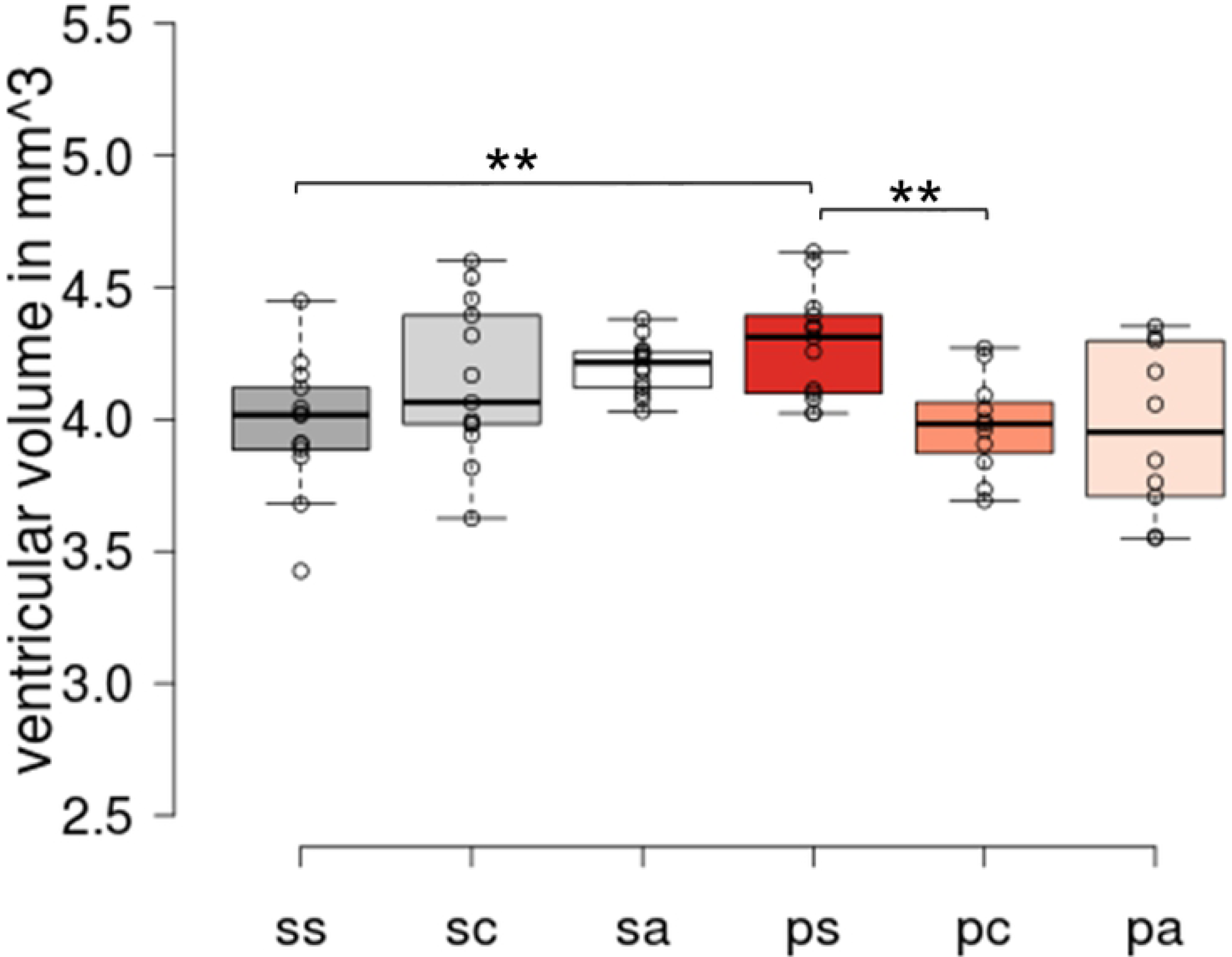
Performance evaluation on treatment data. Lateral ventricle volume estimation in rat MIS model from a study of preventive treatment with transcranial direct current stimulation (8) shows validation of prior reported findings using automated platform method. Center lines show the medians; box limits indicate the 25th and 75th percentiles as determined by R software; whiskers extend 1.5 times the interquartile range from the 25th and 75th percentiles, individual data points included.

## Discussion

The present work describes the implementation of a single atlas-based segmentation framework for adult rat brain analysis, optimized for lateral ventricle segmentation. Volume estimation with this automated framework was validated against the results of a standard manual method. The output of the analysis platform was further used to estimate volumetric changes caused by neuropsychiatric disease and the potential therapeutic treatments in the MIS rat model of schizophrenia on ex-vivo MRI data.

The enlargement of the lateral ventricles is consistently observed in patients with schizophrenia and relevant established animal models. Such alterations are thought to originate from abnormal prenatal brain development. This association has been especially well documented through the study of rodents following prenatal MIS. Notably, this phenotype difference was observed in both of our independent sample analysis, and with both manual and automated analysis.

In schizophrenia patients further structural brain abnormalities have been detected in in a number of brain regions such as the striatum (44,45), the hippocampus and other medial temporal lobe structures (44,46,47), as well as the cerebellum (48–50), and progressive gray matter (GM) loss is present in parietal, prefrontal and superior temporal cortices (51–54). While we have not investigated this further, recent semi-automated segmentation methods have already shown promising results in estimating changes in striatal, hippocampal and cortical structures in rodent models (Crum et al., 2017).

The use of rodent models in general provides an advantage for the evaluation of temporal changes in disease and repeatability at a systemic level. Animals of different strains combined with several imaging techniques can provide insight regarding behavioral pharmacology, potential therapies, longitudinal studies of diseases in order to measure changes, treatment efficacy and other therapeutic interventions (17,55). Therefore, our results further strengthen the growing field of neuroimaging techniques in preclinical animal research, for which investigations and deductions can be translated towards the respective human conditions (56,57).

Our analysis included ex-vivo MRI data from a study of tDCS application to prevent manifestation of schizophrenia associated deficits in MIA rats (8). tDCS is a brain stimulation method were low electric positive (anodal) or negative (cathodal) current is applied to an area of the brain for the depolarization or hyperpolarization of neurons towards facilitatory or inhibitory behavioral effects respectively (58). Medial prefrontal cortex (mPFC) stimulation, based on established protocols (59) and applied during adolescence, was reported to restrict the emergence of schizophrenia-related behaviors as well as volumetric changes. The previously, manually segmented data sets were now reanalyzed to confirm phenotype and treatment effects at adulthood. We were able to replicate the results with our automated analysis platform, showing a significant difference in lateral ventricle volumes with lower volumes in saline offspring compared to poly I:C offspring. In addition, the automated analysis platform confirmed lower lateral ventricle sizes for poly I:C offspring treated with anodal stimulation but not for poly I:C offspring treated with cathodal stimulation.

Manual segmentation is still frequently used to achieve volumetric measures from neuroimaging techniques due to its robustness, although it has certain disadvantages. It is very time consuming, taking 2-3 hours per animal for a practiced expert, and therefore labor-intensive. Variance in the results is high, with inter- and intra-rater biases, and the need for a better alternative is formidable. The automated method as shown in this paper is accurate, allows repetitiveness and is much less time-consuming. However, automated segmentation still comes with various challenges. They include often low image resolution of rodent brains (smaller voxel sizes in rodents compared to humans due to their brain size), physiological noise and signal-to-noise ratio (57,60). However, animal MRI scanners have higher field strengths (7 - 16.4 T) than human scanners (17,60), providing better contrasts to distinguish brain structures. The quality of MR image contrast influences the quality of extraction, introducing a bias by raters during delineation for low quality images. It should be noted that for well contrasted images, MRI acquisition requires a longer scanning time and large image outputs influence calculations during image analysis. Alternatively, image contrasts agents can be used during scanning to enhance the quality of images while compensating for scanning time (60,61).

In addition, segmentation of small regions of interest can be faulted by introducing a systematic bias, which increases as the region of interest gets smaller (26,62). This has been noted especially for the case of lateral ventricle segmentation. It has also been noted to apply even more so when earlier timepoints are investigated as the brains and corresponding ROIs are much smaller. This systematic bias is inherent in automated segmentation, demanding a critical and cautious adjustment of automated processes to the data at hand. Several steps have been taken in the development of this analysis platform and the preparation of data to reduce a potential systematic bias. In the used registration method, a brain atlas with its corresponding brain mask is used for brain extraction. Therefore, the brain extraction is dependent on the availability of an atlas, its mask and the parameters of registration. The rats’ size, age and the similarity between the reference atlas image and the target image of the dataset must be considered. To aid this, a reference subject can be chosen from the study cohort, manually segmented and used for the automated registration brain extraction methods. In our approach, the *SkullStrip* (37) command tool, a robust automated brain extraction tool, was used, which is dependent on the image resolution. This tool was further optimized by binarizing the skull stripped output for each subject and applying it to the original image to get the best “brain-only” volume image for each subject. These additional steps resulted in a higher correlation in visual inspection, total brain volume and similarity measure to the manual method. These efforts may be partially responsible for achieving results in line with previous work on characterizing volumetric changes in lateral ventricles with the presented platform contrary to the findings of other semi-automated segmentation approaches (26).

Limitations of the presented analysis platform include the still labor-intensive manual extracting of the reference subject and its brain mask used with the optimized skull-strip method. This step, though advisable to remove bias resulting from differences in strain and age when needed, is only carried out once. A within-study template is highly rocommended. Second the image contrast, significant for brain segmentation was low in our data. There is a need for a standard segmentation protocol detailing the process to delineate rodent brains for imaging modalities other than histology. Anderson et al. (63) described a validation of registration parameter choices using high-performance computer clusters. They indicated the computational cost-benefits and temporal performance with a voxel-based analysis. With several images to be analyzed in preclinical studies and the differences observed in between registration metrics there is still a strong need for a comprehensive evaluation framework to inform or guide parameter choices in rodent image analysis. Finally, this automated analysis platform was optimized only for the segmentation of ventricular volumes. Label propagation utilizes the transformation matrix and output of the registration, so multiple labels can be propagated to measure changes in the individual subject. These labels can contain different numbers of structures as well. In the future the presented analysis platform should thus be extended to the measurement of other clinically relevant brain alterations in rodent models.

## Conclusion

The characterization of preclinical rodent models of neurological and psychiatric diseases with several imaging modalities is challenging with an increased need for accuracy, reputability, efficiency and reduced human effort except for quality control in the studies. This project thus focused on implementing an automated single atlas-based segmentation framework for rat brains and validate its performance against the results of the standard manual method. The analysis platform integrated several pre-processing and post-processing steps including brain extraction, intensity non-uniformity bias correction, linear and non-linear registrations and label propagation. The resulting analysis platform was used exemplarily with ex-vivo MR images of a MIS schizophrenia rat model. It successfully detected lateral ventricle volumetric changes for phenotypes in independent samples and following treatments, establishing a basis for utilizing anatomical biomarkers in the rat to investigate disease progression or treatment efficiency.

## Acknowledgements

We thank Kristin Wogan for excellent technical assistance and Susanne Müller for profound support on MRI studies. Also the authors gratefully acknowledge the GWK support for funding this project by providing computing time through the Center for Information Services and HPC (ZIH) at TU Dresden on the HRSK-II.

## Supporting information

**S1 Appendix. Supplement**

## References

1. Yoo AJ, Sheth KN, Kimberly WT, Chaudhry ZA, Elm JJ, Jacobson S, et al. Validating Imaging Biomarkers of Cerebral Edema in Patients With Severe Ischemic Stroke. Journal of Stroke and Cerebrovascular Diseases. 2013 Aug;22(6):742–9.

2. Westeneng H-J, Verstraete E, Walhout R, Schmidt R, Hendrikse J, Veldink JH, et al. Subcortical structures in amyotrophic lateral sclerosis. Neurobiology of Aging. 2015 Feb 1;36(2):1075–82.

3. Kato A, Ibara S, Maruyama Y, Terahara M. Relationship between enlargement of the lateral ventricle and periventricular leukomalacia in infants. Journal of Obstetrics and Gynaecology Research. 2010;36(5):984–90.

4. Nestor SM, Rupsingh R, Borrie M, Smith M, Accomazzi V, Wells JL, et al. Ventricular enlargement as a possible measure of Alzheimer’s disease progression validated using the Alzheimer’s disease neuroimaging initiative database. Brain. 2008 Sep;131(9):2443–54.

5. Lewis MM, Smith AB, Styner M, Gu H, Poole R, Zhu Hongtu, et al. Asymmetrical lateral ventricular enlargement in Parkinson’s disease. Eur J Neurol. 2009 Apr;16(4):475–81.

6. van Erp TGM, Hibar DP, Rasmussen JM, Glahn DC, Pearlson GD, Andreassen OA, et al. Subcortical brain volume abnormalities in 2028 individuals with schizophrenia and 2540 healthy controls via the ENIGMA consortium. Mol Psychiatry. 2016 Apr;21(4):547–53.

7. Mata I, Perez-Iglesias R, Roiz-Santiañez R, Tordesillas-Gutierrez D, Gonzalez-Mandly A, Berja A, et al. Additive effect of NRG1 and DISC1 genes on lateral ventricle enlargement in first episode schizophrenia. NeuroImage. 2010 Nov 15;53(3):1016–22.

8. Hadar R, Winter R, Edemann-Callesen H, Wieske F, Habelt B, Khadka N, et al. Prevention of schizophrenia deficits via non-invasive adolescent frontal cortex stimulation in rats. Molecular Psychiatry. 2019 Jan 28;1.

9. Fatemi SH, Reutiman TJ, Folsom TD, Huang H, Oishi K, Mori S, et al. Maternal infection leads to abnormal gene regulation and brain atrophy in mouse offspring: Implications for genesis of neurodevelopmental disorders. Schizophrenia Research. 2008 Feb;99(1–3):56–70.

10. Piontkewitz Y, Arad M, Weiner I. Abnormal Trajectories of Neurodevelopment and Behavior Following In Utero Insult in the Rat. Biological Psychiatry. 2011 Nov;70(9):842–51.

11. Segal Y, Segal L, Blumenfeld-Katzir T, Sasson E, Poliansky V, Loeb E, et al. The Effect of Electromagnetic Field Treatment on Recovery from Ischemic Stroke in a Rat Stroke Model: Clinical, Imaging, and Pathological Findings. Stroke Res Treat [Internet]. 2016 [cited 2019 Jun 3];2016. Available from: https://www.ncbi.nlm.nih.gov/pmc/articles/PMC4753339/

12. Bataveljić D, Djogo N, Župunski L, Bajić A, Nicaise C, Pochet R, et al. Live monitoring of brain damage in the rat model of amyotrophic lateral sclerosis.: 7.

13. Andjus PR, Bataveljić D, Vanhoutte G, Mitrecic D, Pizzolante F, Djogo N, et al. In Vivo Morphological Changes in Animal Models of Amyotrophic Lateral Sclerosis and Alzheimer’s-Like Disease: MRI Approach. The Anatomical Record. 2009;292(12):1882–92.

14. Descloux C, Ginet V, Rummel C, Truttmann AC, Puyal J. Enhanced autophagy contributes to excitotoxic lesions in a rat model of preterm brain injury. Cell Death Dis [Internet]. 2018 Aug 28 [cited 2019 Jun 3];9(9). Available from: https://www.ncbi.nlm.nih.gov/pmc/articles/PMC6113308/

15. Vernon AC, Crum WR, Johansson SM, Modo M. Evolution of Extra-Nigral Damage Predicts Behavioural Deficits in a Rat Proteasome Inhibitor Model of Parkinson’s Disease. PLoS One [Internet]. 2011 Feb 25 [cited 2019 Jun 3];6(2). Available from: https://www.ncbi.nlm.nih.gov/pmc/articles/PMC3045435/

16. Finlay CJ, Duty S, Vernon AC. Brain Morphometry and the Neurobiology of Levodopa-Induced Dyskinesias: Current Knowledge and Future Potential for Translational Pre-Clinical Neuroimaging Studies. Front Neurol [Internet]. 2014 Jun 12 [cited 2020 Mar 31];5. Available from: https://www.ncbi.nlm.nih.gov/pmc/articles/PMC4053925/

17. Strome EM, Doudet DJ. Animal models of neurodegenerative disease: insights from in vivo imaging studies. Mol Imaging Biol. 2007 Aug;9(4):186–95.

18. Ashburner J, Friston KJ. Voxel-Based Morphometry—The Methods. NeuroImage. 2000 Jun 1;11(6):805–21.

19. Clarkson MJ, Cardoso MJ, Ridgway GR, Modat M, Leung KK, Rohrer JD, et al. A comparison of voxel and surface based cortical thickness estimation methods. NeuroImage. 2011 Aug 1;57(3):856–65.

20. Das SR, Avants BB, Grossman M, Gee JC. Registration based cortical thickness measurement. NeuroImage. 2009 Apr 15;45(3):867–79.

21. Shen K, Fripp J, Mériaudeau F, Chételat G, Salvado O, Bourgeat P. Detecting global and local hippocampal shape changes in Alzheimer’s disease using statistical shape models. NeuroImage. 2012 Feb 1;59(3):2155–66.

22. Budin F, Hoogstoel M, Reynolds P, Grauer M, O’Leary-Moore SK, Oguz I. Fully automated rodent brain MR image processing pipeline on a Midas server: from acquired images to region-based statistics. Frontiers in Neuroinformatics [Internet]. 2013 [cited 2019 Nov 18];7. Available from: http://journal.frontiersin.org/article/10.3389/fninf.2013.00015/abstract

23. Ma Y, Smith D, Hof PR, Foerster B, Hamilton S, Blackband SJ, et al. In Vivo 3D Digital Atlas Database of the Adult C57BL/6J Mouse Brain by Magnetic Resonance Microscopy. Front Neuroanat [Internet]. 2008 Apr 17 [cited 2020 Mar 31];2. Available from: https://www.ncbi.nlm.nih.gov/pmc/articles/PMC2525925/

24. Jorge Cardoso M, Leung K, Modat M, Keihaninejad S, Cash D, Barnes J, et al. STEPS: Similarity and Truth Estimation for Propagated Segmentations and its application to hippocampal segmentation and brain parcelation. Medical Image Analysis. 2013 Aug 1;17(6):671–84.

25. Nestor SM, Gibson E, Gao F-Q, Kiss A, Black SE. A Direct Morphometric Comparison of Five Labeling Protocols for Multi-Atlas Driven Automatic Segmentation of the Hippocampus in Alzheimer’s Disease. Neuroimage. 2013 Feb 1;0:50–70.

26. Crum WR, Sawiak SJ, Chege W, Cooper JD, Williams SCR, Vernon AC. Evolution of structural abnormalities in the rat brain following in utero exposure to maternal immune activation: A longitudinal in vivo MRI study. Brain Behav Immun. 2017 Jul;63:50–9.

27. Norris FC, Modat M, Cleary JO, Price AN, McCue K, Scambler PJ, et al. Segmentation propagation using a 3D embryo atlas for high-throughput MRI phenotyping: Comparison and validation with manual segmentation. Magnetic Resonance in Medicine. 2013;69(3):877–83.

28. Meyer U, Feldon J, Schedlowski M, Yee BK. Towards an immuno-precipitated neurodevelopmental animal model of schizophrenia. Neuroscience & Biobehavioral Reviews. 2005 Jan;29(6):913–47.

29. Meyer U, Feldon J, Dammann O. Schizophrenia and Autism: Both Shared and Disorder-Specific Pathogenesis Via Perinatal Inflammation?: Pediatric Research. 2011 May;69(5 Part 2):26R–33R.

30. Piontkewitz Y, Assaf Y, Weiner I. Clozapine Administration in Adolescence Prevents Postpubertal Emergence of Brain Structural Pathology in an Animal Model of Schizophrenia. Biological Psychiatry. 2009 Dec 1;66(11):1038–46.

31. Hadar R, Soto-Montenegro ML, Götz T, Wieske F, Sohr R, Desco M, et al. Using a maternal immune stimulation model of schizophrenia to study behavioral and neurobiological alterations over the developmental course. Schizophrenia Research. 2015 Aug 1;166(1):238–47.

32. Zuckerman L, Rehavi M, Nachman R, Weiner I. Immune Activation During Pregnancy in Rats Leads to a PostPubertal Emergence of Disrupted Latent Inhibition, Dopaminergic Hyperfunction and Altered Limbic Morphology in the Offspring: A Novel Neurodevelopmental Model of Schizophrenia. Neuropsychopharmacology. 2003 Oct;28(10):1778–89.

33. Paxinos G, Watson C. The rat brain in stereotaxic coordinates. 6th ed. Amsterdam□ Boston: Academic Press/Elsevier; 2007. 1 p.

34. Ma D, Cardoso MJ, Modat M, Powell N, Wells J, Holmes H, et al. Automatic Structural Parcellation of Mouse Brain MRI Using Multi-Atlas Label Fusion. PLOS ONE. 2014 Jan 27;9(1):e86576.

35. Badea A, Ali-Sharief AA, Johnson GA. Morphometric analysis of the C57BL/6J mouse brain. NeuroImage. 2007 Sep 1;37(3):683–93.

36. Sled JG, Zijdenbos AP, Evans AC. A nonparametric method for automatic correction of intensity nonuniformity in MRI data. IEEE Trans Med Imaging. 1998 Feb;17(1):87–97.

37. Boyes RG, Gunter JL, Frost C, Janke AL, Yeatman T, Hill DLG, et al. Intensity non-uniformity correction using N3 on 3-T scanners with multichannel phased array coils. Neuroimage. 2008 Feb 15;39(4):1752–62.

38. Kochunov P, Lancaster JL, Thompson P, Woods R, Mazziotta J, Hardies J, et al. Regional spatial normalization: toward an optimal target. J Comput Assist Tomogr. 2001 Oct;25(5):805–16.

39. Kovacević N, Henderson JT, Chan E, Lifshitz N, Bishop J, Evans AC, et al. A three-dimensional MRI atlas of the mouse brain with estimates of the average and variability. Cereb Cortex. 2005 May;15(5):639–45.

40. Benveniste H, Blackband S. MR microscopy and high resolution small animal MRI: applications in neuroscience research. Prog Neurobiol. 2002 Aug;67(5):393–420.

41. Lee J, Jomier J, Aylward S, Tyszka M, Moy S, Lauder J, et al. Evaluation of Atlas based Mouse Brain Segmentation. Proc SPIE. 2009 Feb 1;7259:725943–9.

42. Ochsner KN, Beer JS, Robertson ER, Cooper JC, Gabrieli JDE, Kihsltrom JF, et al. The neural correlates of direct and reflected self-knowledge. Neuroimage. 2005 Dec;28(4):797–814.

43. Fujiwara H, Yassin W, Murai T. Neuroimaging studies of social cognition in schizophrenia. Psychiatry and Clinical Neurosciences. 2015;69(5):259–67.

44. Bogerts B, Meertz E, Schönfeldt-Bausch R. Basal ganglia and limbic system pathology in schizophrenia. A morphometric study of brain volume and shrinkage. Arch Gen Psychiatry. 1985 Aug;42(8):784–91.

45. Buchabaum MS. Frontal Lobes, Basal Ganglia, Temporal Lobes—Three Sites for Schizophrenia? Schizophr Bull. 1990 Jan 1;16(3):377–8.

46. Benes FM, Sorensen I, Bird ED. Reduced Neuronal Size in Posterior Hippocampus of Schizophrenic Patients. Schizophrenia Bulletin. 1991 Jan 1;17(4):597–608.

47. Nelson MD, Saykin AJ, Flashman LA, Riordan HJ. Hippocampal volume reduction in schizophrenia as assessed by magnetic resonance imaging: a meta-analytic study. Arch Gen Psychiatry. 1998 May;55(5):433–40.

48. DeLisi LE, Sakuma M, Tew W, Kushner M, Hoff AL, Grimson R. Schizophrenia as a chronic active brain process: a study of progressive brain structural change subsequent to the onset of schizophrenia. Psychiatry Res. 1997 Jul 4;74(3):129–40.

49. Volz H-P, Gaser C, Sauer H. Supporting evidence for the model of cognitive dysmetria in schizophrenia — a structural magnetic resonance imaging study using deformation-based morphometry. Schizophrenia Research. 2000 Nov 30;46(1):45–56.

50. Ichimiya T, Okubo Y, Suhara T, Sudo Y. Reduced volume of the cerebellar vermis in neurolepticnaive schizophrenia. Biological Psychiatry. 2001 Jan 1;49(1):20–7.

51. Kuperberg GR, Broome MR, McGuire PK, David AS, Eddy M, Ozawa F, et al. Regionally localized thinning of the cerebral cortex in schizophrenia. Arch Gen Psychiatry. 2003 Sep;60(9):878–88.

52. White T, Andreasen NC, Nopoulos P, Magnotta V. Gyrification abnormalities in childhood- and adolescent-onset schizophrenia. Biol Psychiatry. 2003 Aug 15;54(4):418–26.

53. Wiegand LC, Warfield SK, Levitt JJ, Hirayasu Y, Salisbury DF, Heckers S, et al. Prefrontal Cortical Thickness in First-Episode Psychosis: A Magnetic Resonance Imaging Study. Biol Psychiatry. 2004 Jan 15;55(2):131–40.

54. Narr KL, Bilder RM, Toga AW, Woods RP, Rex DE, Szeszko PR, et al. Mapping cortical thickness and gray matter concentration in first episode schizophrenia. Cereb Cortex. 2005 Jun;15(6):708–19.

55. Borsook D, Becerra L. CNS Animal fMRI imaging in Pain and Analgesia. Neurosci Biobehav Rev. 2011 Apr;35(5):1125–43.

56. McIntosh AL, Gormley S, Tozzi L, Frodl T, Harkin A. Recent Advances in Translational Magnetic Resonance Imaging in Animal Models of Stress and Depression. Front Cell Neurosci [Internet]. 2017 May 24 [cited 2020 Mar 31];11. Available from: https://www.ncbi.nlm.nih.gov/pmc/articles/PMC5442179/

57. Kalavathi P, Prasath VBS. Methods on Skull Stripping of MRI Head Scan Images—a Review. J Digit Imaging. 2016 Jun;29(3):365–79.

58. Nitsche MA, Cohen LG, Wassermann EM, Priori A, Lang N, Antal A, et al. Transcranial direct current stimulation: State of the art 2008. Brain Stimul. 2008 Jul;1(3):206–23.

59. Thair H, Holloway AL, Newport R, Smith AD. Transcranial Direct Current Stimulation (tDCS): A Beginner’s Guide for Design and Implementation. Front Neurosci [Internet]. 2017 Nov 22 [cited 2020 Mar 31];11. Available from: https://www.ncbi.nlm.nih.gov/pmc/articles/PMC5702643/

60. Balaban RS, Hampshire VA. Challenges in small animal noninvasive imaging. ILAR J. 2001;42(3):248–62.

61. Hoyer C, Gass N, Weber-Fahr W, Sartorius A. Advantages and Challenges of Small Animal Magnetic Resonance Imaging as a Translational Tool. NPS. 2014;69(4):187–201.

62. Lau JC, Lerch JP, Sled JG, Henkelman RM, Evans AC, Bedell BJ. Longitudinal neuroanatomical changes determined by deformation-based morphometry in a mouse model of Alzheimer’s disease. NeuroImage. 2008 Aug 1;42(1):19–27.

63. Anderson RJ, Cook JJ, Delpratt N, Nouls JC, Gu B, McNamara JO, et al. Small Animal Multivariate Brain Analysis (SAMBA) - a High Throughput Pipeline with a Validation Framework. Neuroinformatics. 2019;17(3):451–72.

64. Tustison NJ, Avants BB, Cook PA, Zheng Y, Egan A, Yushkevich PA, et al. N4ITK: Improved N3 Bias Correction. IEEE Transactions on Medical Imaging. 2010 Jun;29(6):1310–20.

65. Delora A, Gonzales A, Medina CS, Mitchell A, Mohed AF, Jacobs RE, et al. A simple rapid process for semi-automated brain extraction from magnetic resonance images of the whole mouse head. Journal of Neuroscience Methods. 2016 Jan 15;257:185–93.

